## Supplemental Figure S1 for "A Ribosome Interaction Surface Sensitive to mRNA GCN Periodicity"

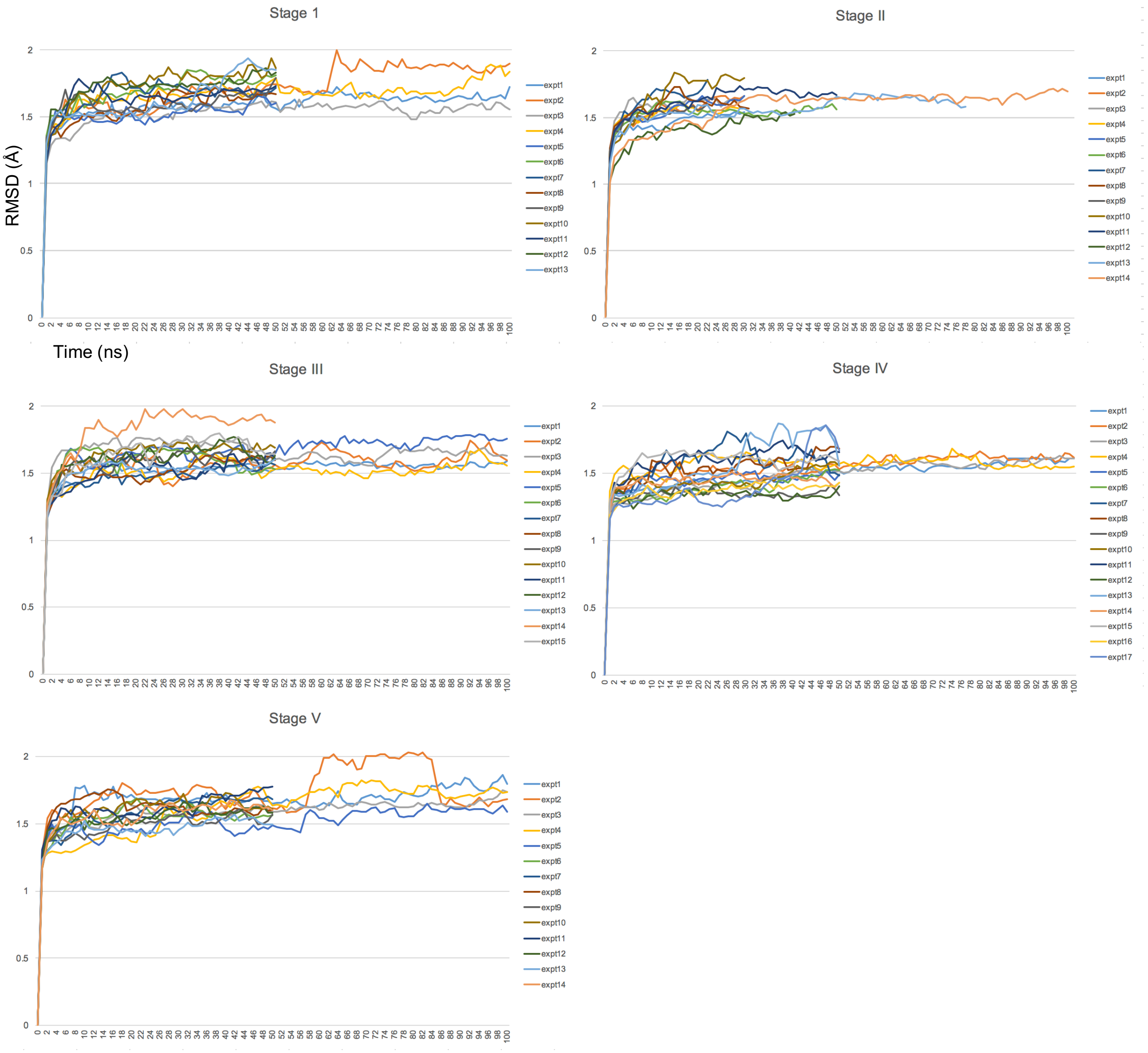


**Figure S1.** RMSD profiles for MD replicates. For each translocation stage, multiple MD replicates were run starting with independent heating, assignment of velocities and equilibration. RMSD distances (Å) were collected from 100 frames per ns and the average for each ns was graphed. RMSD stabilized below 2 Å within 10-15 ns and further analysis of these MD trajectories was performed starting at 15 ns. Stage II MD runs sometimes terminated early, and for some of these cases, we performed energy minimization of structures before termination which showed local temperature elevations near the restrained onion shell. These subsystems of the ribosome, composed of RNA and protein, are considered to represent preferred intermediate structures of the ribosome during translocation.
