## Supplemental Figure S2 for "A Ribosome Interaction Surface Sensitive to mRNA GCN Periodicity"

### H-bonds over time (ns)

#### Stage I

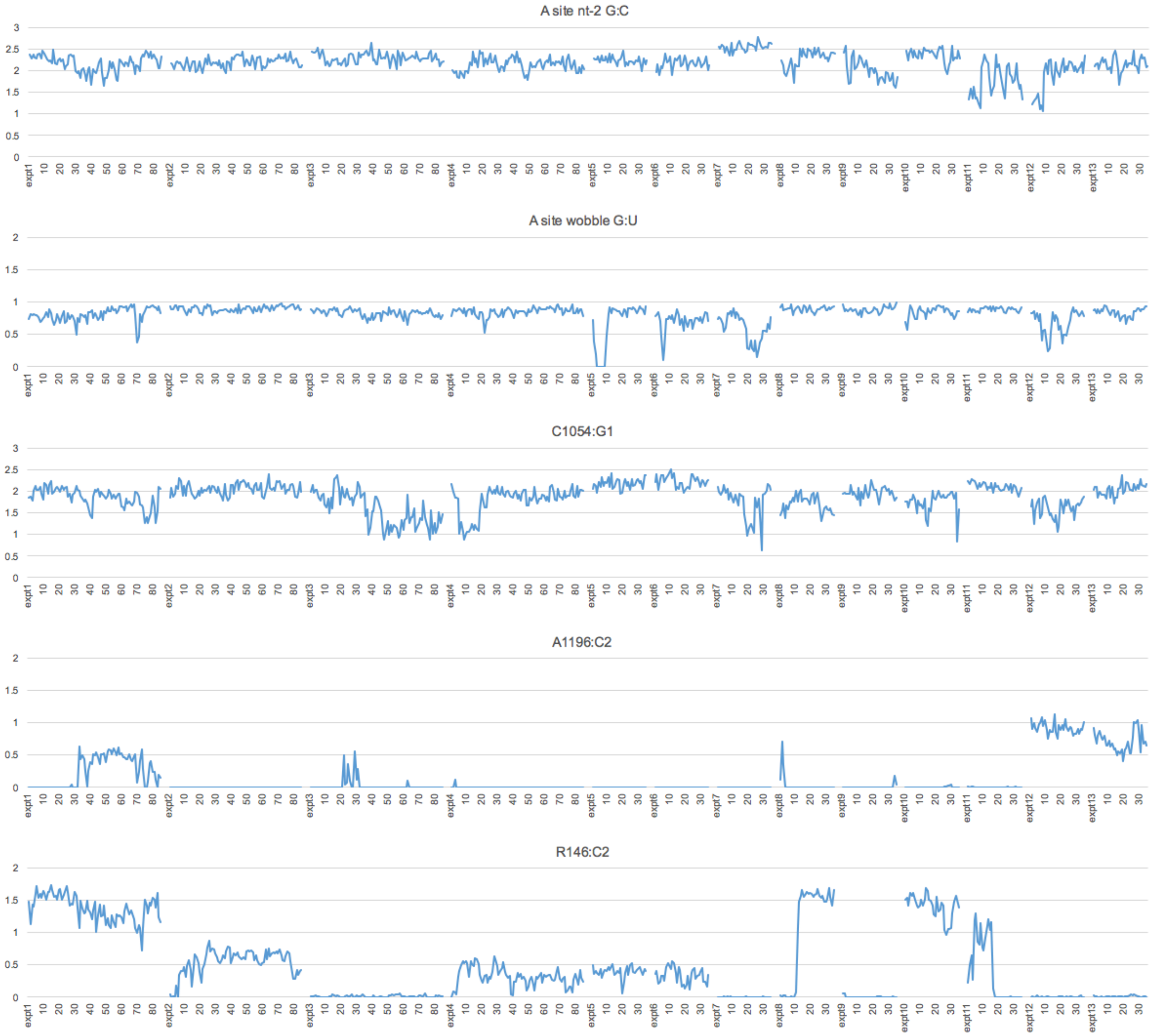

H-bonds over time (ns) cont.

#### Stage II

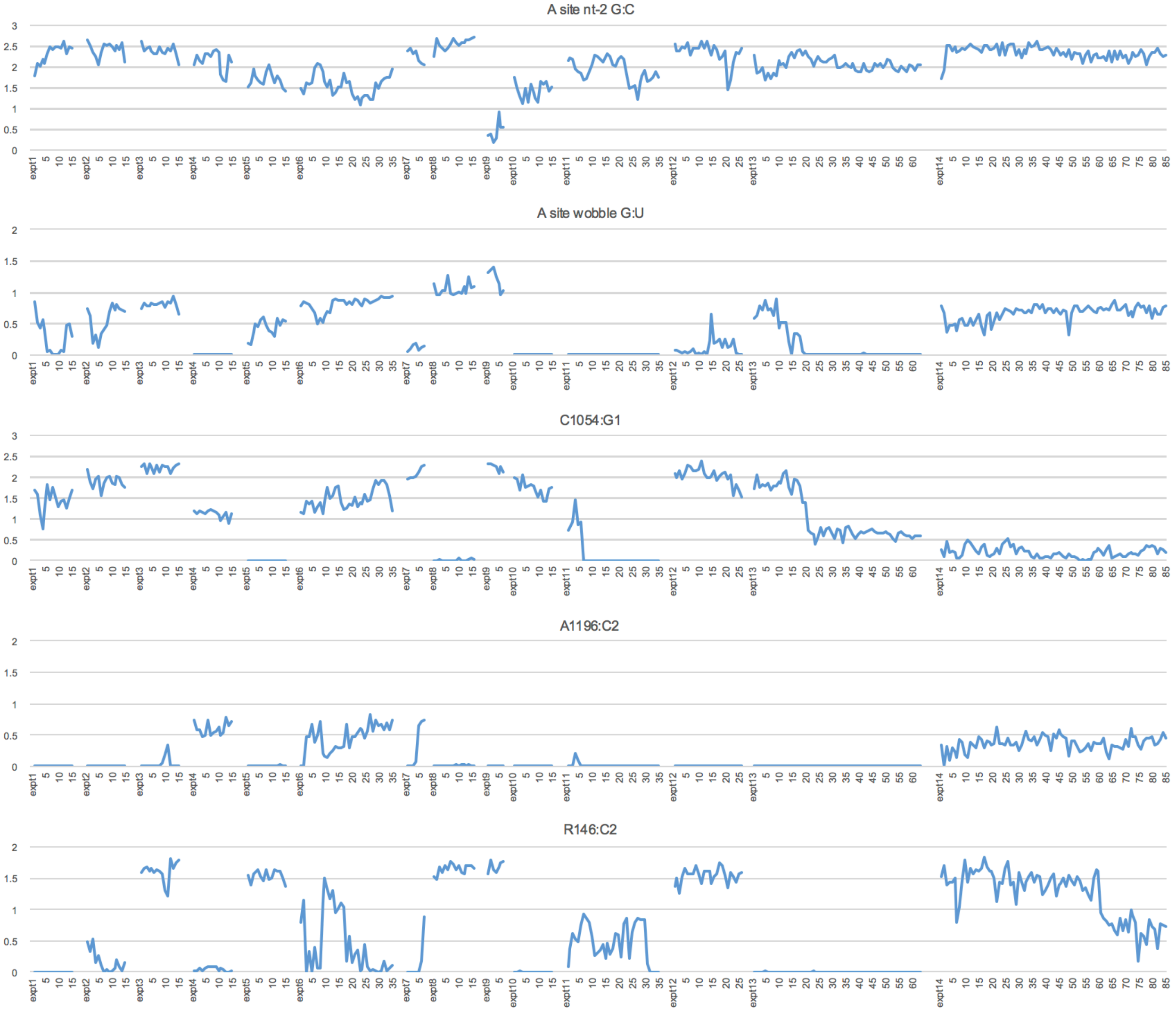

H-bonds over time (ns) cont.

#### Stage III

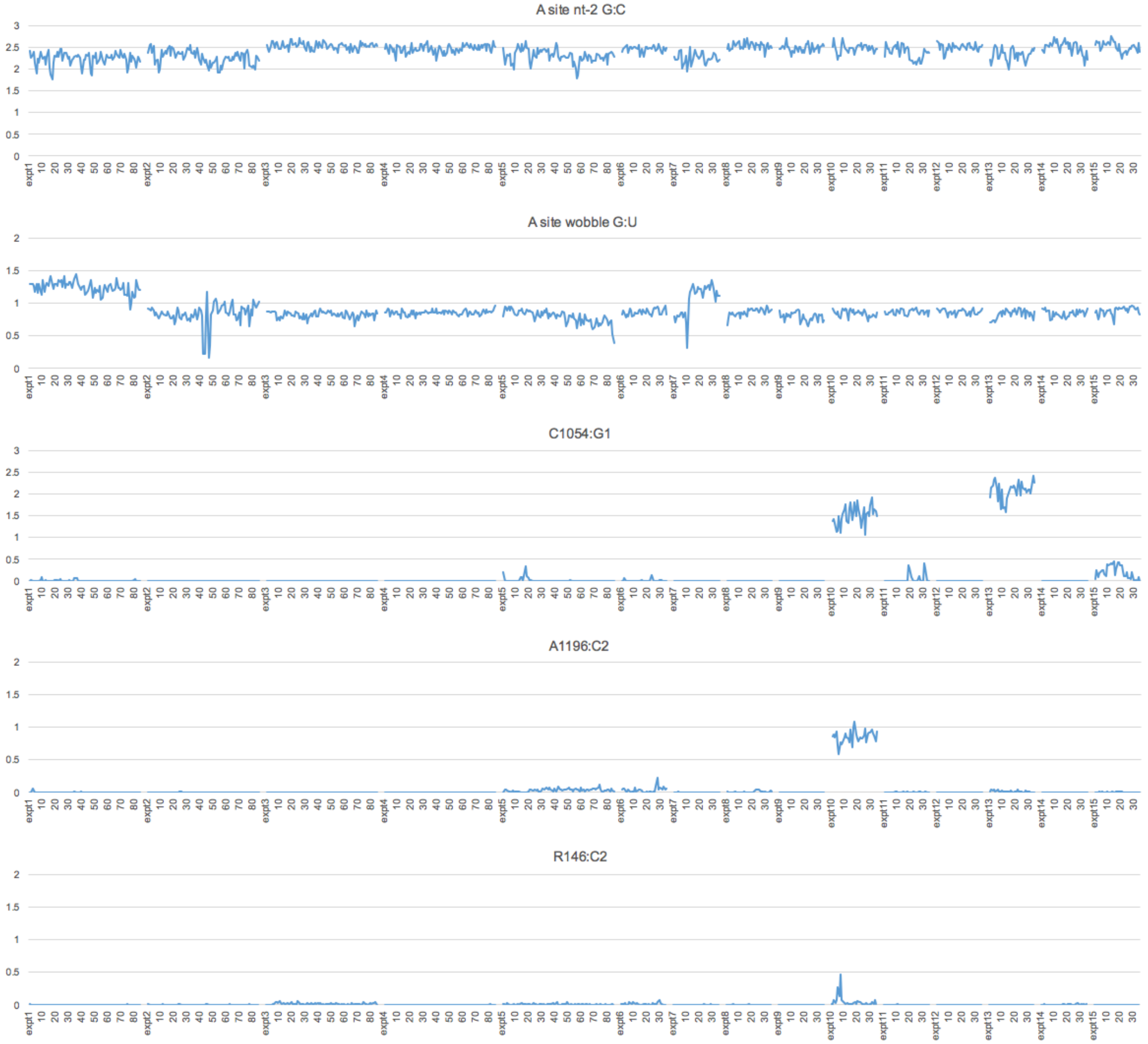

Figure S2

H-bonds over time (ns) cont.

#### Stage IV

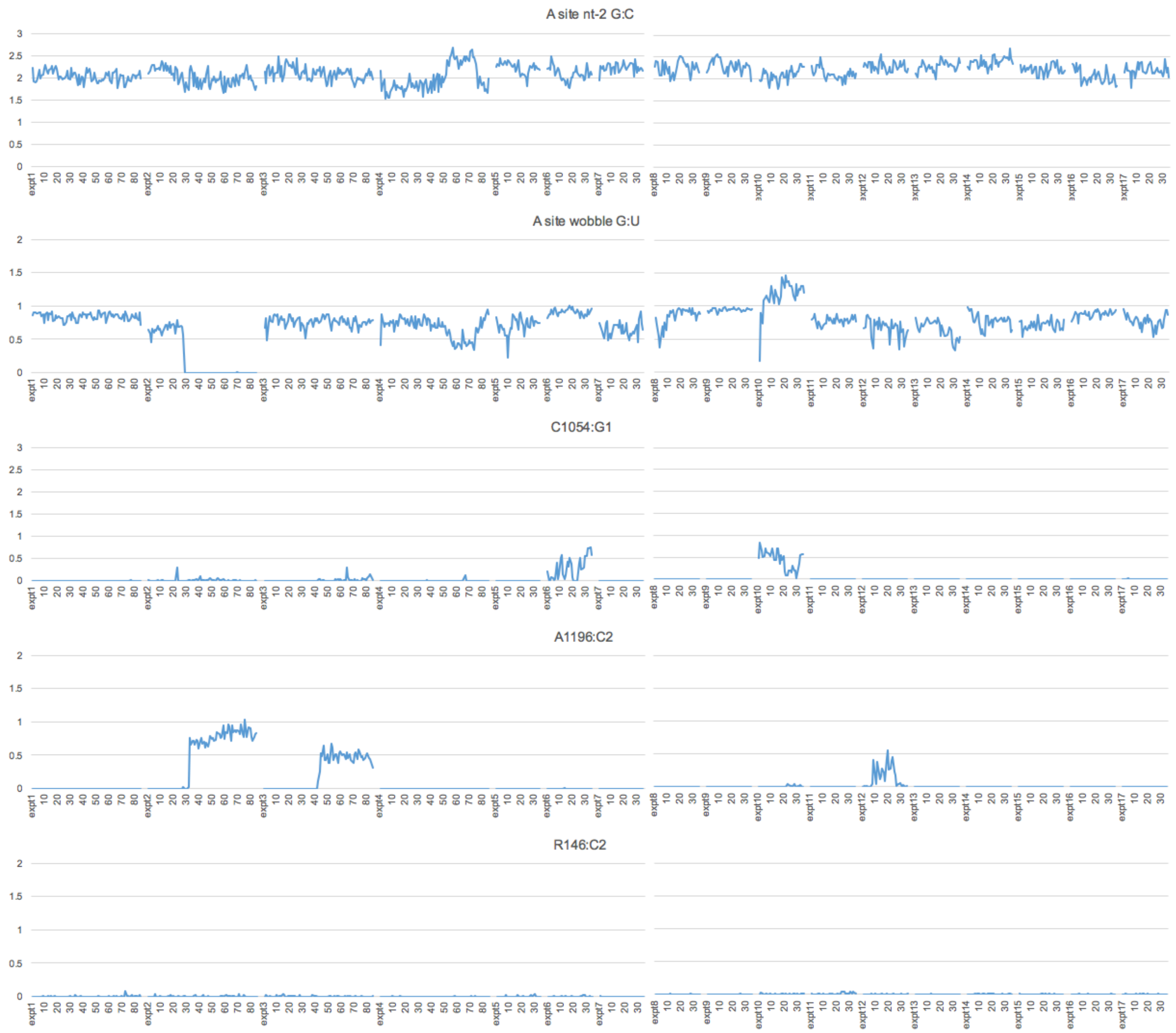

H-bonds over time (ns) cont.

#### Stage V

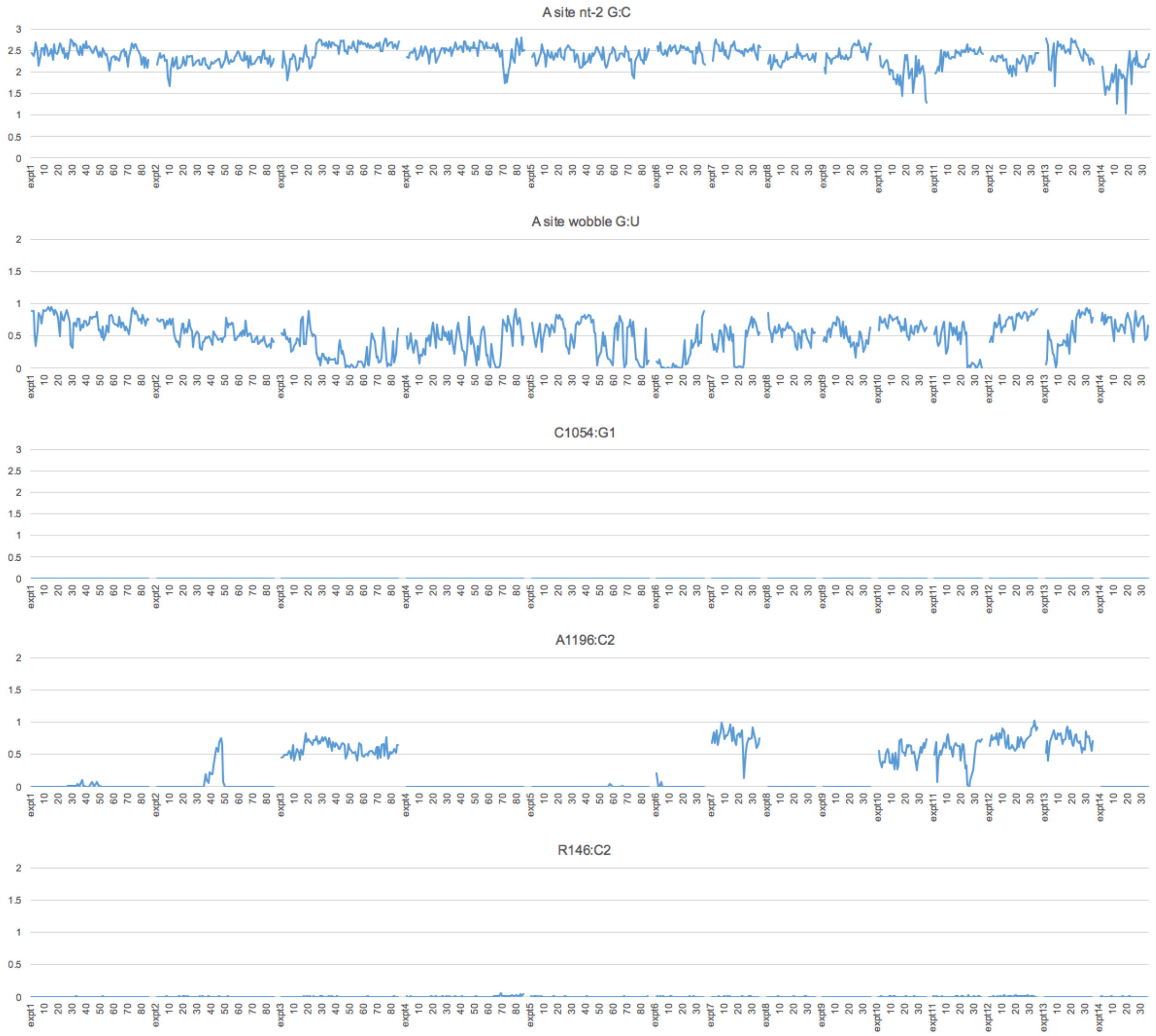


**Figure S2.** Numbers of H-bonds in MD replicate experiments. For each translocation stage (stages I through V), multiple independent MD experiments were run starting with different heat, random assignment of velocities, and equilibration. Numbers of H-bonds are between: A site nt-2 G:C; A site wobble base G:C; C1054:G1 of +1 codon; A1196 Hoogsteen edge:C2 Watson-Crick edge of +1 codon; R146 guanidinium group:C2 Watson-Crick edge. 100 frames were collected per ns of MD and the average number of H-bonds in each ns was plotted.
