## Supplemental Figure S3 for "A Ribosome Interaction Surface Sensitive to mRNA GCN Periodicity"

### Stacking distance (Å) over time (ns)

### Stage I

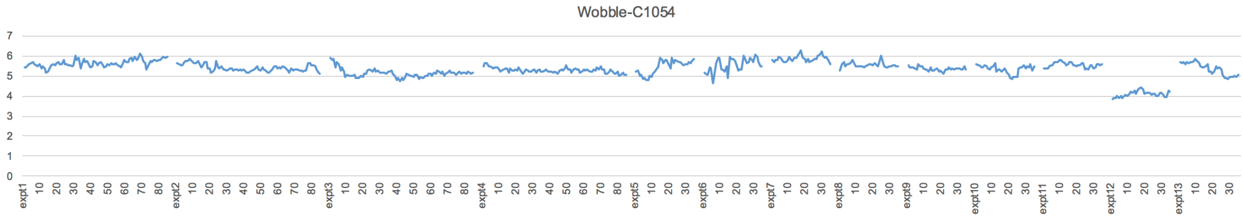


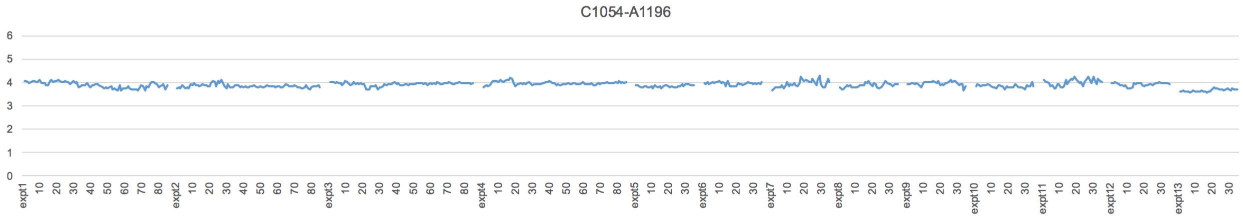


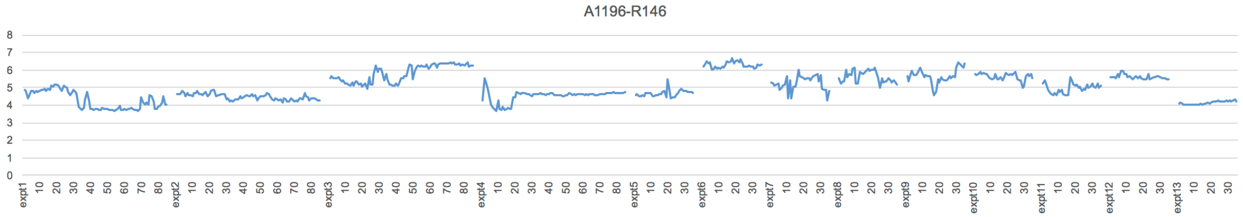


### Stage II

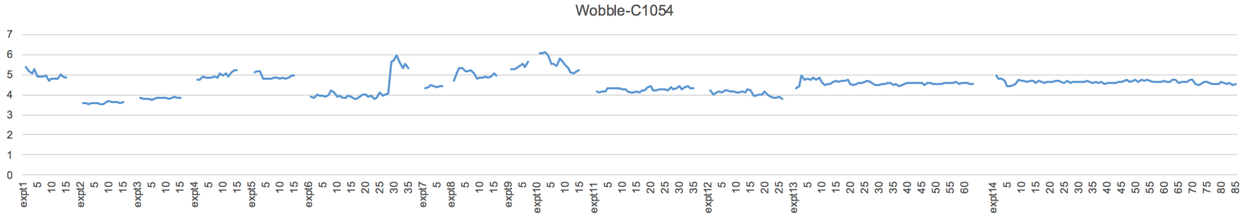


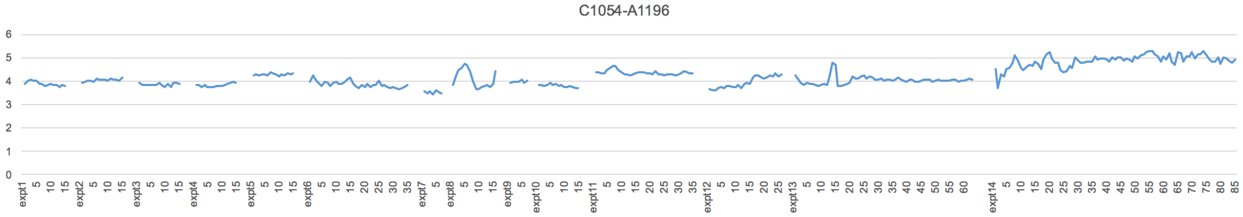


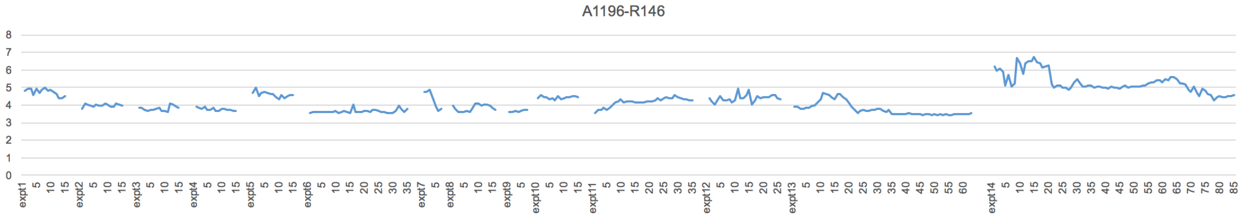

Stacking distance (Å) over time (ns) cont.

### Stage III

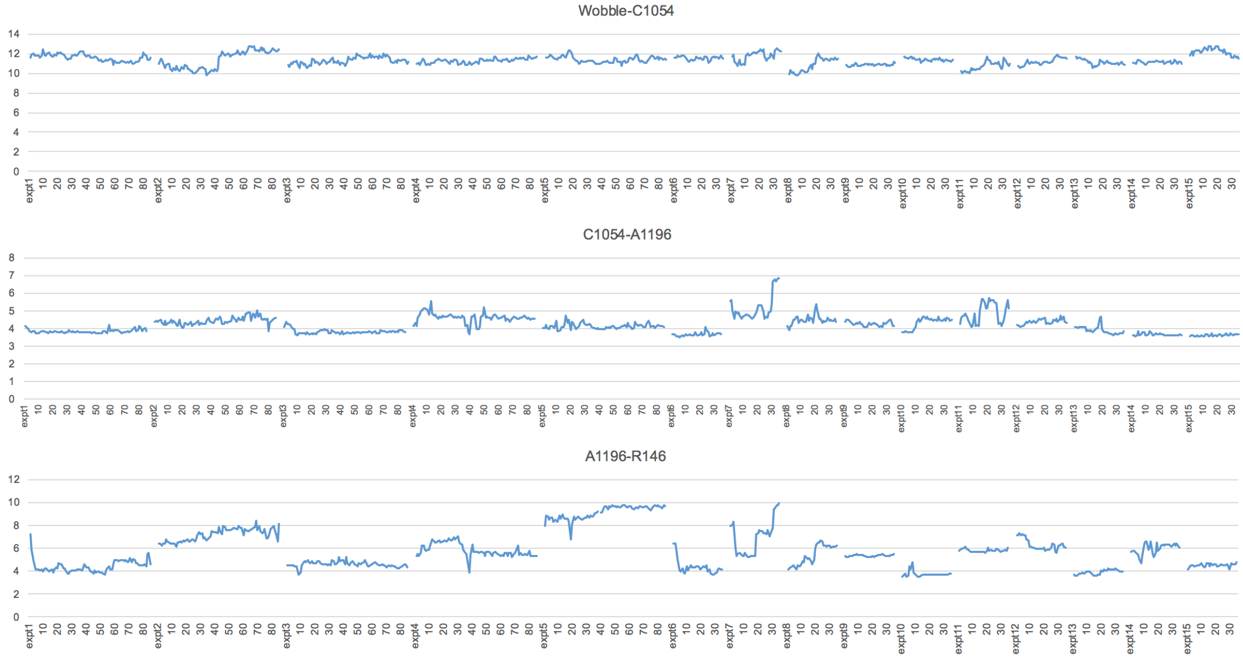


### Stage IV

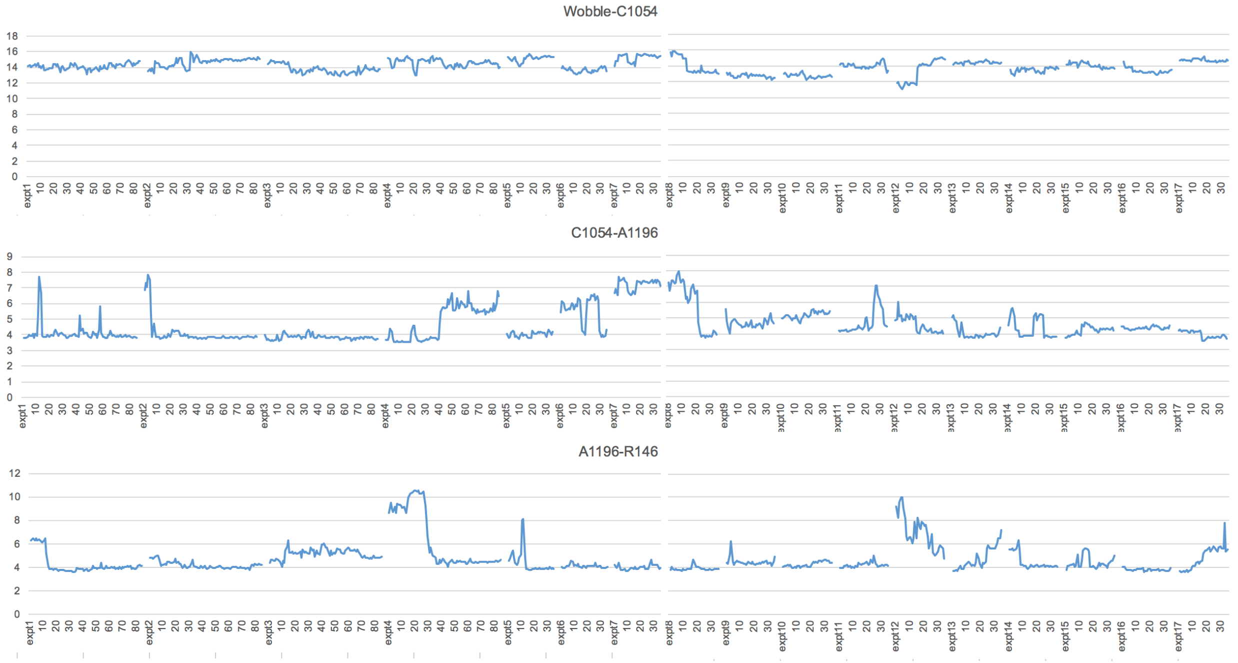

Stacking distance (Å) over time (ns) cont.

Stage V

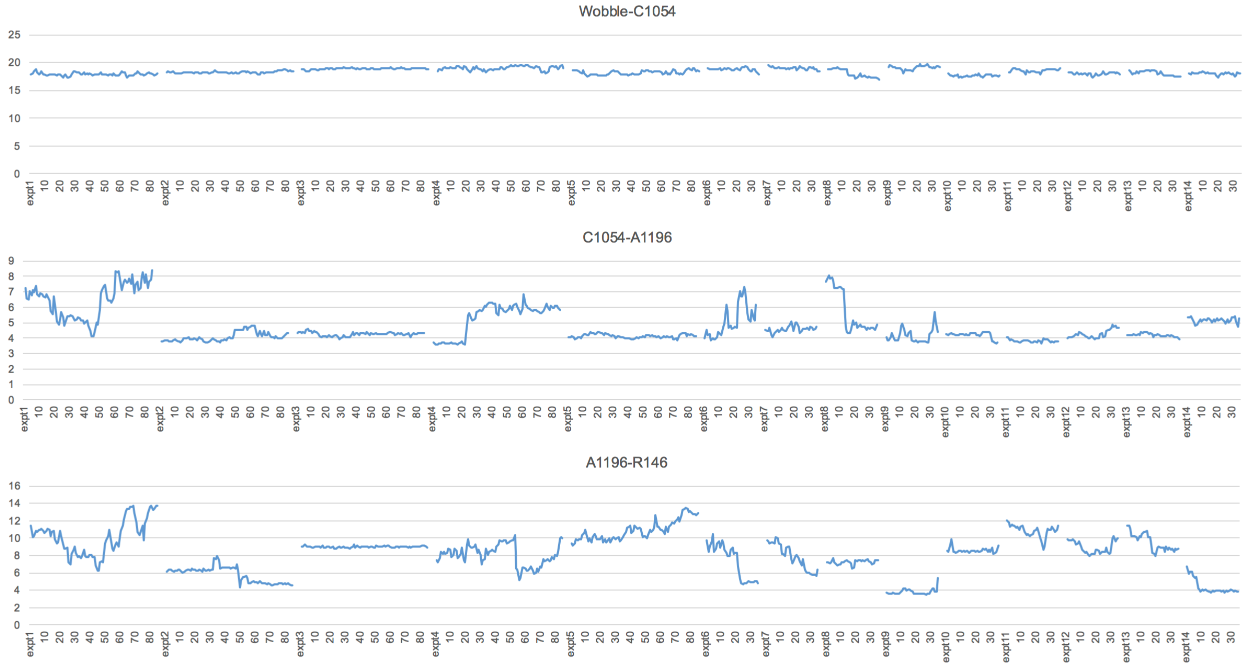


**Figure S3.** Stacking distances in MD replicate experiments. For each translocation stage (stages I through V), multiple independent MD experiments were run starting with different heat, random assignment of velocities, and equilibration. Distances (Å) are between the centers of masses of the bases: A site wobble base-C1054; C1054-A1196; A1196-R146 guanidinium group. Frames were pooled in 1-ns bins. 100 frames were collected per ns of MD and the average distance for each ns was plotted.
