## Supplemental Figure S4 for "A Ribosome Interaction Surface Sensitive to mRNA GCN Periodicity"

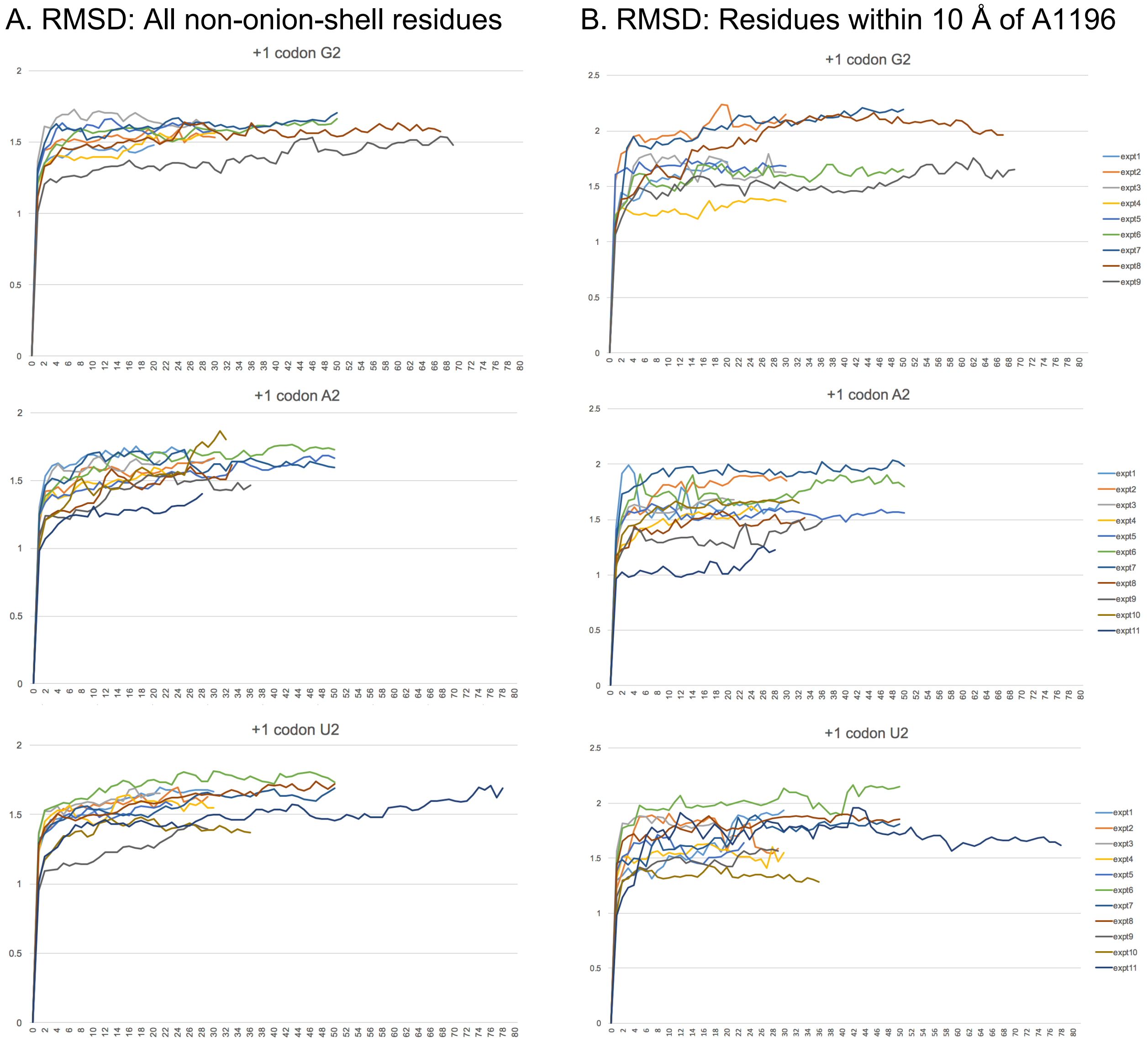


**Figure S4.** RMSD profiles for +1 codon nt-2 substitutions. C at position 2 of the +1 codon (C2) was replaced with G, A or U (G2, A2, U2) in the translocation stage II structure. (A) RMSD profiles for backbone atoms of all residues except those in restrained onion shell. RMSD profiles for multiple independent MD runs stabilized below 2 Å within 10 to 15 ns. The MD runs were analyzed starting at 15 ns. The RMSD profiles commence after 3 ns of equilibration during which backbone atoms of all residues were restrained at 20 kcal/mol Å$^2$. (B) RMSD profiles for backbone atoms of 19 residues located within 10 Å of A1196. MD runs stabilized below 2.5 Å within 10 to 15 ns.
