## Supplemental Figure S5 for "A Ribosome Interaction Surface Sensitive to mRNA GCN Periodicity"

C1054

```
RDN18-1       1244 AGCTCTTTCTTGATTTTGTGGGTGGTGGTGCATGGCCGTTCTTAGTTGGTGGAGTGATTT  1303 Budding yeast
AJ272181.1    1229 ...............C.A....................................C.....  1288 Bread wheat
NR_133559.1   1332 ..........C..A.C.A.............................C............  1391 Drosophila
6G4S_2        1301 ..........C....CC.....................................C.....  1360 Human
JQ247698.1    1303 ..........C....CC.....................................C.....  1362 Mouse
2ZKQ_A        1026 ..........C....CC.....................................C.....  1085 Dog
6P4H_2        1301 ..........C....CC.....................................C.....  1360 Rabbit
AB779691.1    1221 .........A.....C.A..........................................  1280 Plasmodium
5XXU_2        1239 ...............C.A..........................................  1298 Toxoplasma
MK395374.1    1690 T.T.......C....C.T..AA................C.T..G..C.............  1749 Trypanosoma
MK454732.1    1215 ...............C.T..........................................  1274 Tetrahymena
NR_024570.1   1030                 ..A.ACA....C........T..CG.C..C.C...TT....AA.  1073 E. coli
NR_037066.1   1020                 .AGCACA....C...........CG.C..C.C...CC....GG.  1063 T. thermophilus
```

A1196

```
RDN18-1       1408    GGAAGTTTGAGGCAATAACAGGTCTGTGATGCCCTTA  1444  Budding yeast
AJ272181.1    1395    .....................................  1431  Bread wheat
NR_133559.1   1605       ..A.....-.........................  1637  Drosophila
6G4S_2        1474       ..A.....-.........................  1506  Human
JQ247698.1    1476       ..A.....-.........................  1508  Mouse
2ZKQ_A        1168       ..A.....-.........................  1200  Dog
6P4H_2        1474       ..A.....-.........................  1506  Rabbit
AB779691.1    1384    .....................................  1420  Plasmodium
5XXU_2        1404    .....................................  1440  Toxoplasma
MN215474.1    1730     ...A...TG.....C.G..............       1760  Trypanosoma
MK454732.1    1378    ........A.........................C..  1414  Tetrahymena
NR_024570.1   1167    .....G.G.G.ATG.CGT..A...ATCATG.......  1203  E. coli
NR_037066.1   1156    .....GAG.G.A.G.CGT.T....A.CATG.......  1192  T. thermophilus
```

R146

```
YNL178W 121  GVVRYVMESGAKGCEVVVSGKLRAARAKAMKFADGFLIHSGQPVNDFIDTATRHVLMRQG  180 Budding yeast
5IT9_D  119  .................I..........S...................E.......L...  178 K. lactis yeast
3J60_D  110  ..L.F...........I.......Q...S...K..YM.S......EY..A.V....L...  169 Bread wheat
3J38_D  123  ..L..I.................GQ...S...V..LM....D.C..YVE.......L...  182 Drosophila
3J3A_D  121  ..L.FI.................GQ...S...V..LM....D...YYV...V....L...  180 Human
2ZKQ_c  121  ..L.FI.................GQ...S...V..LM....D...YYV...V....L...  180 Dog
5K0Y_g  121  ..L.FI.................GQ...S...V..LM....D...YYV...V....L...  180 Rabbit
3J7A_D  122  ..L.HI..........I.......Q...S...R..Y..ST.E.SKR.VN....SAQLK..  181 Plasmodium
5XXU_D  123  ..L.HI..................Q...S...K..Y..ST.E.SKM.V.Q.I.S.QL...  182 Toxoplasma
3ZEY_X  120  .II.............TIG..IKGQ...S.T.R..YM.K..TAHKH.V......CHL.A.  179 Trypanosoma
2XZM_C  125   IIKS.IQD......III....KQQ...T...KQ.YM.CT...K..Y..V.V...FFK..  183 Tetrahymena
6AZ1_C  120  .II.............T.G..IKGQ...S.T.R..YM.K..TAHKS.V.S.C..CY..A.  179 Leishmania
5XYI_D  122  SII.F..DA..R....AI.....G...STS.YKE.YMVK..DVTKQ.VTQ.VG.IP.K.A  181 Trichomonas
1PNS_C  137    .QR........AK.I...RIGG.EQARTEW.                             167 E. coli
3FIC_C  137    .QR........AK.I...RIGG.EQARTEW.                             167 T. thermophilus
```

**Figure S5.** 16S/18S rRNA C1054, A1196 and ribosomal protein S3 R146 are conserved. Alignments of genomic sequences show that C1054 and A1196 (red arrows) are well conserved in eukaryotes and prokaryotes although A1196 is sometimes T1196 (U1196; e.g. *Thermus thermophilus*). R146 (arrow) is well conserved in eukaryotes, but not prokaryotes. Yeast sequences were used as queries in BLASTN or BLASTP analysis creating alignments to which bacterial sequences were added. Sequence identities are illustrated with dots.
