## Supplemental Table S1 for "A Ribosome Interaction Surface Sensitive to mRNA GCN Periodicity"

| Chain | 5JUP Numbering* | 5JUP Restrained | Subsystem Numbering | Subsystem Restrained |
|---|---|---|---|---|
| A 18S rRNA | 1-31 | 1-5, 11-17, 22-31 | 1-31 | 1-5, 11-17, 22-31 |
| A 18S rRNA | 547-600 | 547-548, 550, 554-556, 588-600 | 32-85 | 32-33, 35, 39-41, 73-85 |
| A 18S rRNA | 1108-1113 | 1108-1113 | 86-91 | 86-91 |
| A 18S rRNA | 1133-1140 | 1133-1136, 1139-1140 | 92-99 | 92-95, 98-99 |
| A 18S rRNA | 1269-1279 | 1269-1270, 1277-1279 | 100-110 | 100-101, 108-110 |
| A 18S rRNA | 1424-1431 | 1424-1425, 1429-1431 | 111-118 | 111-112, 116-118 |
| A 18S rRNA | 1438-1442 | 1438-1442 | 119-123 | 119-123 |
| A 18S rRNA | 1629-1649 | 1629-1631, 1635-1637, 1644-1649 | 124-144 | 124-126, 130-132, 139-144 |
| A 18S rRNA | 1751-1763 | 1751-1752, 1759-1763 | 145-157 | 145-146, 153-157 |
| A 18S rRNA | 1780-1782 | 1780-1782 | 158-160 | 158-160 |
| B 25S rRNA | 2255-2258 | 2255-2256 | 161-164 | 161-162 |
| EC mRNA (IRES) | 6903-6909 | 6903, 6909 | 165-171 | 165, 171 |
| EC mRNA (IRES) | 6947-6958 | | 172-183 | |
| UB uS12 (yeast S23) | 53-145 | 53-56, 73-84, 96-111, 121-134, 138-145 | 184-276 | 184-187, 204-215, 227-242, 252-265, 269-276 |
| ZA uS5 (yeast S2) | 85-98 | 85-98 | 277-290 | 277-290 |
| ZA uS5 (yeast S2) | 193-211 | 193-211 | 291-309 | 291-309 |
| BC eS30 (yeast S30) | 2-25 | 2-4, 22-25 | 310-333 | 310-312, 330-333 |
| BC eS30 (yeast S30) | 45-61 | 45-54, 61 | 334-350 | 334-343, 350 |
| AB uS3 (Yeast S3) | 111-117 | 111-117 | 351-357 | 351-357 |
| AB uS3 (Yeast S3) | 136-153 | 136-138, 150-153 | 358-375 | 358-360, 372-375 |
| AB uS3 (Yeast S3) | 176-180 | 176-177, 179-180 | 376-380 | 376-377, 379-380 |
| DC yeast eEF2 | 578-588 | 578-588 | 381-391 | 381-391 |
| DC yeast eEF2 | 608-711 | 608-660, 667-695, 710-711 | 392-495 | 392-444, 451-479, 494-495 |

*Same residue numbering for 5JUO, 5JUS, 5JUT, and 5JUU.
