## Supplemental Table S2 for "A Ribosome Interaction Surface Sensitive to mRNA GCN Periodicity"

| energy minimization round | harmonic restraint (kcal/mol Å$^2$) | steepest descent steps | conjugate gradient steps |
|---|---|---|---|
| 1 | 100 | 2500 | 17500 |
| 2 | 75 | 2500 | 7500 |
| 3 | 65 | 2500 | 2500 |
| 4 | 55 | 2500 | 500 |
| 5 | 45 | 2500 | 500 |
| 6 | 30 | 2000 | 0 |
| 7 | 20 | 2000 | 0 |
| 8 | 15 | 2000 | 0 |
| 9 | 10 | 2000 | 0 |
| 10 | 5 | 2000 | 0 |
| 11 | 1 | 2000 | 0 |
